# Live imaging of delamination in *Drosophila* shows epithelial cell motility and invasiveness are independently regulated

**DOI:** 10.1101/2021.05.08.443270

**Authors:** Mikiko Inaki, Kenji Matsuno

**Affiliations:** Department of Biological Sciences, Graduate School of Science, Osaka University, 1-1 Machikaneyama-cho, Toyonaka, Osaka 560-0043, Japan

**Keywords:** delamination, cell migration, locomotion, invasion, motility

## Abstract

Delaminating cells undergo complex, precisely regulated changes in cell-cell adhesion, motility, polarity, invasiveness, and other cellular properties. Delamination occurs during development and in pathogenic conditions such as cancer metastasis. We analyzed the requirements for epithelial delamination in *Drosophila* ovary border cells, which detach from the structured epithelial layer and begin to migrate collectively. We used live imaging to examine cellular dynamics, particularly epithelial cells’ acquisition of motility and invasiveness, in delamination-defective mutants during the time period in which delamination occurs in the wild-type ovary. We found that border cells in *slow border cells* (*slbo*), a delamination-defective mutant, lacked invasive cellular protrusions but acquired motility, while JAK/STAT-inhibited border cells lost both invasiveness and motility. Our results indicate that invasiveness and motility, which are cooperatively required for delamination, are regulated independently. Our reconstruction experiments also showed that motility is not a prerequisite for acquiring invasiveness.

**Summary statement:** Live imaging of epithelial delamination reveals that cell motility and invasiveness, both of which are required for delamination, are regulated independently and that motility is not a prerequisite for invasiveness.

## Introduction

The delamination of epithelial cells, which normally form a layered structure, occurs when the cells become motile and detach from the organized cell layer. In the developing embryo, delamination allows cells to leave the epithelial layer and move where they are needed. Disrupting this precisely regulated process results in morphological abnormalities, cancer metastasis, and other pathological conditions (Stuelten et al., 2018; Thiery et al., 2009). Delamination occurs in *Drosophila* gastrulation when the endodermal layer invaginates; epithelial cells detach from the layer and convert into mesenchymal cells to form the mesoderm (Ko and Martin, 2020; Thiery et al., 2009). In vertebrates, neural crest cells delaminate from the dorsal neural tube and migrate to form bone, neurons, glia, and other mesodermal cells (Gouignard et al., 2018; Szabó and Mayor, 2018). Delamination also occurs during cancer metastasis, in which tumor cells lose adhesion, digest epithelial-tissue basal membranes, and invade the body cavity (Stuelten et al., 2018; Thiery et al., 2009).

Although delamination is also called epithelial-to-mesenchymal transition (EMT), a term that reflects the dynamic changes in cell polarity, adhesion, and cytoskeletal and membrane structures (Campbell and Casanova, 2016; Nieto et al., 2016; Yang et al., 2020), the delaminated cells often retain apicobasal polarity and other epithelial cell characteristics. The requirements for epithelial delamination remain elusive. To define these requirements, we analyzed the delamination of *Drosophila* border cells, which derive from the follicle-cell layer that surrounds germline cells—that is, the oocyte and supporting nurse cells in the developing egg chambers of the *Drosophila* ovary (Montell, 2006). The cells at the ends of the follicular layer, called pole cells, have distinct properties. Anterior pole cells secrete signals that activate JAK/STAT signaling in neighboring follicle cells (Silver and Montell, 2001). Follicle cells that receive JAK/STAT signaling start expressing *slow border cells* (*slbo*), which encodes a C/EBP transcription factor, and become motile border cells (Montell et al., 1992). After delaminating from the follicular layer, border cells migrate collectively toward the oocyte, where they are essential for fertilization. Although this collective migratory process has been studied extensively as a model for cancer metastasis (Bianco et al., 2007; Cai et al., 2014; Cliffe et al., 2017; Dai et al., 2020; Inaki et al., 2012; Wang et al., 2010; Yang et al., 2012), little is known about the process by which border cells delaminate. Border cells retain apicobasal polarity and cell-cell adhesion during delamination (De Graeve et al., 2012; Pinheiro and Montell, 2004; Szafranski and Goode, 2004; Wang et al., 2018). Actin regulators such as Profilin and Fascin have been implicated in delamination (Ghiglione et al., 2018; Lamb et al., 2020). However, the changes in cellular properties prerequisite to delamination need further clarification.

We used live imaging to analyze the dynamic cellular changes involved in delamination. Live imaging studies of delamination in various systems, including EMT during neural crest cell development and at gastrulation, have described morphological features in delaminating cells and the involvement of actomyosin contractility (Clay and Halloran, 2013; Liu et al., 2017; Ramkumar et al., 2016; Saykali et al., 2019). However, the hierarchy of gene regulation has not been analyzed. Here, we analyzed border cell delamination at single-cell resolution through a live imaging system that allowed us to genetically dissect and reconstruct the complex cellular processes of delamination. We found that *slbo*-mutant border cells, which have been considered non-motile (Montell et al., 1992), acquire basic motility. This motility is lost in JAK/STAT-defective border cells. We also found that forced Slbo expression in JAK/STAT-defective border cells rescued the cells’ ability to form invasive protrusions, but did not rescue motility or delamination. These results indicate that epithelial cells acquire motility and invasiveness separately, and that motility is not a prerequisite for invasiveness.

## Results

### Locomotion is impaired upon JAK/STAT inhibition, but not in *slbo* mutants

Delaminating border cells in the *Drosophila* ovary undergo a series of complex cellular changes (Fig. 1A). After the cells are specified and distinguished from other follicle cells, they start to send out small extensions (Fig. 1A). Their affinity to other follicle cells changes, and the shape of the cell cluster becomes more rounded (Fig. 1A2-6). The cluster sends out front extensions, mostly single, in the direction the cluster will migrate (Fig. 1A3-6). The extensions progressively elongate and contract, eventually forming a large protrusion, and the body of the cluster moves toward the protrusion and finally detaches from the epithelial layer (Fig. 2A).

**Fig. 1.**
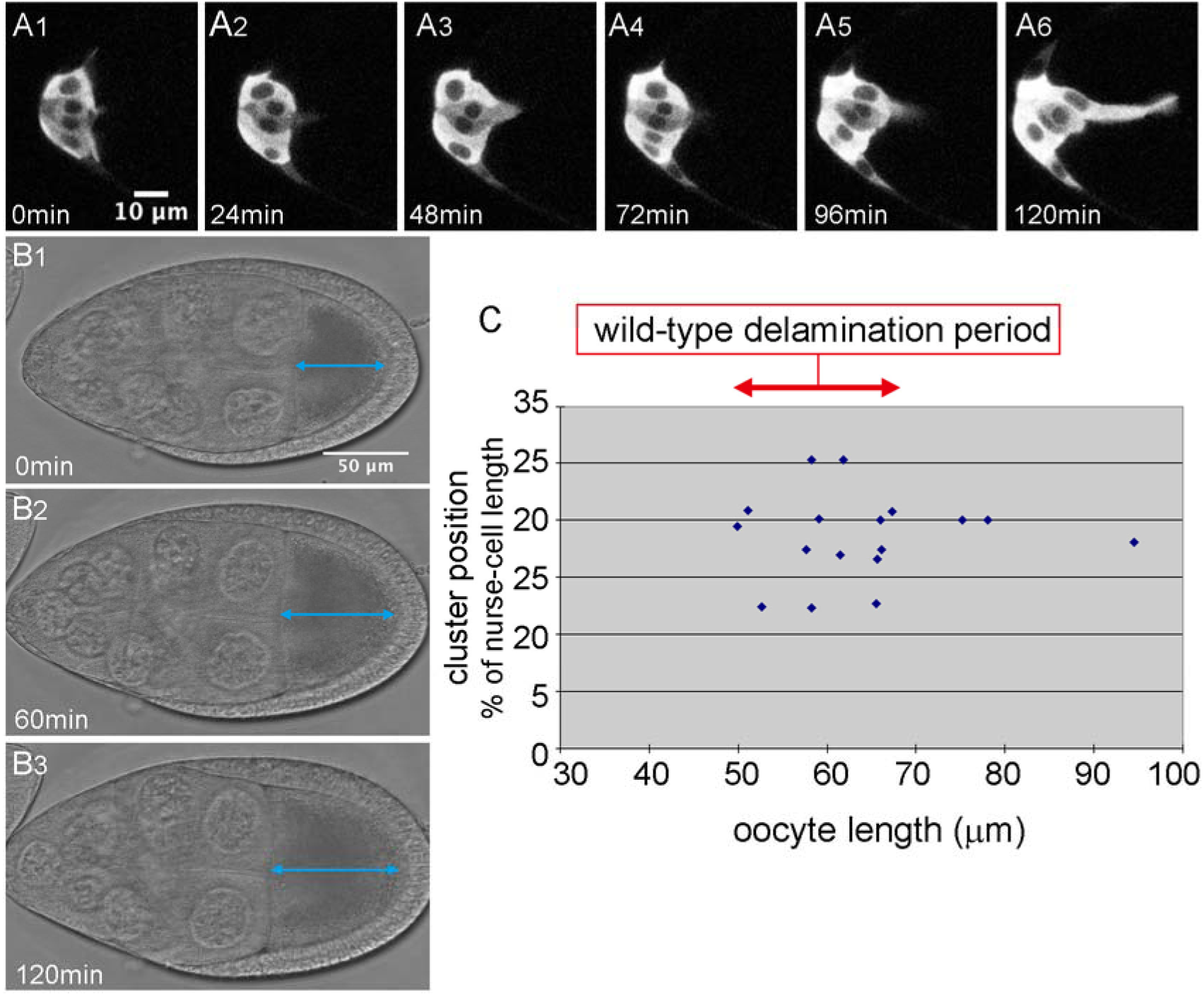
Delamination is a complex cellular process. (A) A time series showing the delamination process in wild-type *Drosophila* border cells. (B) A time series of transmission images of the egg chamber during delamination. Blue arrows indicate oocyte length, which increased with time. (C) Oocyte length at the delamination point was determined from 17 movies of wild-type border cells. We defined an oocyte length of 50-70 μm as a marker of the wild-type delamination period. Elapsed time from the start of the movie is shown at lower left. For all images, anterior is left.

**Fig. 2.**
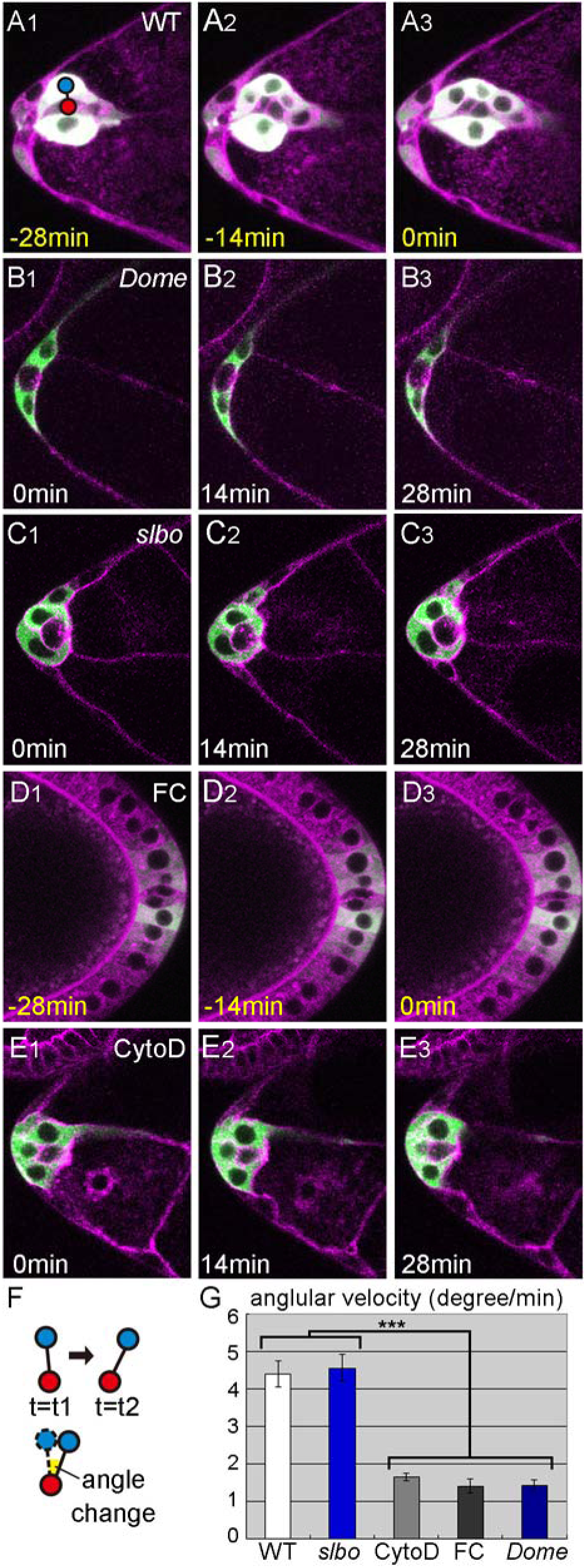
Locomotion is impaired upon JAK/STAT inhibition in wild-type but not in *slbo*-mutant border cells. (A-E) Single Z slices from movies show the delamination of wild-type (A), DN-Dome expressing (B), *slbo*-mutant (C), and CytochalasinD-treated (D) border cells and a movie of posterior follicle cells (E). Red and blue circles indicate the cluster center and a border cell nucleus, respectively. (F) A schematic showing how cell movement was quantified by measuring the change in the angle of the border-cell nucleus (blue circle) relative to the cluster center (red circle). (G) The angular velocity of wild-type (WT), DN-Dome-expressing (*Dome*), *slbo*-mutant (*slbo*), and CytochalasinD-treated (CytoD) border cells relative to the cluster center, and the angular velocity of posterior follicle cells relative to the posterior pole cell center (FC). 12≦ number of cells (n) ≦24, 6≦number of egg chambers (N) ≦12. Error bars indicate SEM. ***, p<0.001. Time shown at the lower left indicates elapsed time from the start of the movie (white) and time prior to the delamination point (yellow). For all images, anterior is left.

We investigated the cellular properties required for epithelial cell delamination by live imaging of border cells in delamination-defective mutants. To compare the behavior of wild-type and delamination-defective cells during the period in which wild-type border cells become migratory, it was necessary to define developmental egg chamber stages. Border cells are thought to delaminate around egg-chamber stage 8 to early stage 9, but there are no objective markers for these stages. Follicle-cell retraction has been used to define stage 9, but it is hard to judge exactly when retraction starts. To obtain internal markers for egg chamber development, we measured several parameters, including the length and width of the oocyte and egg chamber, from movies of delamination in wild-type cells. We chose oocyte length as a suitable marker for egg-chamber stages because it consistently increases over time and appears to be a major contributor to egg chamber growth (Fig. 1B1-3). Based on our analysis of 19 movies of wild-type delamination, we defined the delamination time as the point when the entire cluster, including trailing cells, has detached from the follicular layer. In movies of wild-type cells, 85% of delamination events occurred when the oocyte was 50–70 microns long (Fig. 1C). Thus, we used oocyte length as a marker of the wild-type delamination period, corresponding to an oocyte length of 50–70 microns (Fig. 1C).

Movies of wild-type cells were analyzed for the 30-min period prior to the delamination point. Movies of delamination-defective mutants were analyzed for the 30-min period starting with the first frame corresponding to the wild-type delamination period, assessed by oocyte length. During this period, wild-type border cells form a round cluster that jiggles and forms an extension in the direction of future migration (Fig. 2A1-3, Movie 1). In delamination-defective mutants, we first analyzed movies of egg chambers with JAK/STAT inhibition, which causes border cell differentiation to fail (Silver and Montell, 2001). Follicle cells next to anterior pole cells use a receptor named Domeless (Dome) to receive ligands from the anterior pole cell and activate JAK/STAT signaling. To inhibit JAK/STAT signaling, we expressed a dominant-negative (DN) form of Dome receptor lacking the cytoplasmic domain (Dome^ΔCYT^) specifically in follicle cells, where border cells are differentiated, using *slbo*-gal4 along with UAS-10×GFP as a cellular marker (Poukkula et al., 2011; Rørth et al., 1998; Silver and Montell, 2001). The JAK/STAT-inhibited border cells appeared to retain an epithelial morphology with apicobasal polarity, as do other follicle cells, and were not motile (Fig. 2B1-3, Movie 2). Next, we examined border cells in a mutant for *slbo*, a downstream target of JAK/STAT signaling in border cells (Silver and Montell, 2001). We found that border cells in homozygotes for the hypomorphic allele *slbo^1310^*, in which border cell migration is almost completely abolished (Montell et al., 1992; Rørth et al., 1998), failed to delaminate during the wild-type delamination period (Fig. 2C). However, their behavior differed from JAK/STAT-inhibited border cells (Fig. 2B,C) in that they formed round clusters in which individual border cells moved back and forth, similar to locomotion (Fig. 2C1-3, Movie 3). We quantified this movement as angular velocity by measuring the movement of border cells relative to the center of the cluster (Fig. 2F). Angular velocity in *slbo*-mutant border cells was similar to wild-type, but was much lower in JAK/STAT-inhibited border cells—similar, in fact, to that of non-motile follicle cells in the posterior end (Fig. 2D,G). We confirmed the loss of cell motility in another allelic combination of *slbo*, *slbo^1310^/slbo^e7b^*, in which border cell migration is completely blocked (Fig. S1, (Montell *et al*., 1992). These results indicate that *slbo*-mutant border cells acquired motility, allowing them to move about, even though they could not invade the egg chamber. To see whether this locomotion in individual cells was an active process involving cytoskeletal forces, we used the inhibitor CytochalasinD to block actin polymerization. CytochalasinD treatment completely blocked border cell delamination (n=13) and cell movement (Fig. 2E,G, Movie 4), indicating that the observed locomotive behavior is an active process and is required for delamination.

### Invasiveness is impaired in *slbo* mutants

Although the locomotive behavior of *slbo*-mutant border cells was similar to that of wild-type border cells, *slbo*-mutant border cell clusters failed to delaminate and had no front extensions (Fig. 3A,B). Wild-type border-cell clusters send extensions toward the direction of future migration (Fig. 3A). These protrusions extend and retract over time, and although a cell with extensions is sometimes replaced by another, a cell with extensions is almost always present during the delamination period (Movie 1). We quantified the front extensions as previously described (Poukkula et al., 2011) with slight modifications. Briefly, we binarized Z-stack images, separated the cluster body and extension by scanning a circle with a diameter slightly larger than that of the cell nucleus, and measured the size (length) and persistence (duration) of the front extensions. We found that extensions formed by wild-type border-cell clusters were up to 18 microns long and persisted through nearly the entire 30-min analysis period; in contrast, *slbo*-mutant border cells had no front extensions (Fig. 3B,E,G,H, Movie 3). We concluded that *slbo*-mutant border cells acquired locomotion (motility) but not invasiveness, which might explain why they failed to delaminate. Inhibiting actin polymerization with CytochalasinD completely blocked the formation (n=9) and maintenance (n=17) of front extensions in wild-type border cells, showing that the extension formation and maintenance are active processes of the actin cytoskeleton. These results indicate that wild-type delaminating border cells acquire motility and invasiveness separately, and that invasiveness is regulated by *slbo* while motility is regulated by other factors downstream of JAK/STAT signaling.

**Fig. 3.**
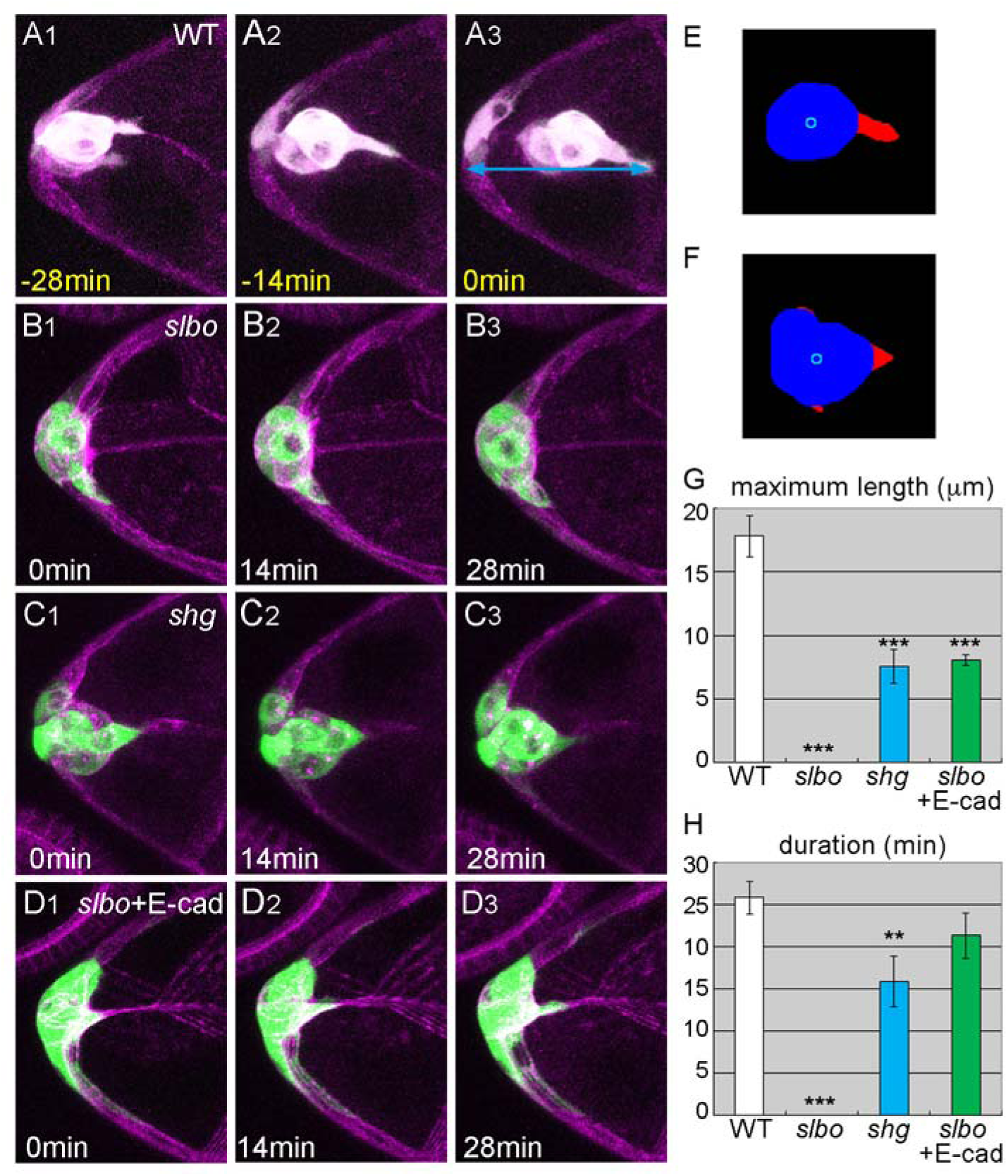
Invasiveness is regulated by *slbo*, partly though *shg*. (A-D) Z stacks from delamination movies of border cells, as follows: wild-type (A), *slbo*-mutant (B), *shg*-mutant (C), and *slbo*-mutant with E-cad expression (D). Light blue arrows indicate the distance (reach) from the anterior end of the egg chamber to the tip of the border-cell extension at delamination (A3). (E,F) The cluster body (blue) and extensions (red) separated by image analysis from the images shown in (A2,C2). Light blue circles: the centroid of the cluster body. (G,H) The maximum length (G) and duration (H) of the front extensions of border-cell clusters, as follows: wild-type (WT), *slbo*-mutant (*slbo*), *shg*-mutant (*shg*), and *slbo*-mutant with E-cad expression (*slbo*+E-cad). 6≦N≦14. Error bars indicate SEM. ***, p<0.001, **, p<0.01. Time shown at the lower left indicates time elapsed from the start of the movie (white) and the time prior to the delamination point (yellow). For all images, anterior is left.

### Invasiveness induced by *slbo* is mediated partly by *shg*

Slbo upregulates gene expression of *shotgun* (*shg*), which encodes E-cadherin (E-cad) in border cells (Mathieu et al., 2007; Tepass et al., 1996). We investigated the *shg* mutant *shg^PB4354^*, in which *shg* upregulation in border cells is specifically suppressed (Mathieu et al., 2007). During the wild-type delamination period, *shg^PB4354^*-mutant border cells did not delaminate but had normal locomotion, as did *slbo*-mutant border cells (Fig. 3C, Fig. S2, Movie 5). Unlike *slbo*-mutant border-cell clusters, *shg^PB4354^*-mutant border-cell clusters had front extensions (Fig. 3C, Movie 5); however, these extensions were much shorter than wild-type in both length and duration (Fig. 3E-H), suggesting that *shg* is required both to form and to maintain invasive extensions. These results explain why *shg^PB4354^*-mutant border cells could not delaminate during the wild-type delamination period. Next, we tested whether the overexpression of E-cad in border cells, which rescues *shg^PB4354^* mutants, could rescue *slbo*-mutant phenotypes (Mathieu et al., 2007). The *slbo*-mutant border cells with forced E-cad expression had front extensions, suggesting that E-cad expression induced by Slbo is responsible for border cell invasiveness (Fig. 3D, Movie 6). However, the front extensions were still significantly shorter than in wild-type border cells (Fig. 3G,H) and E-cad expression did not rescue the impaired delamination phenotype in *slbo* mutants, suggesting that other components downstream of *slbo* are also involved in delamination.

### Invasive protrusions must have sufficient reach to attach across distances

To determine how front extensions contribute to delamination, we examined delamination in guidance-deficient mutants since the extensions must be guided toward the path of future migration. Border cell migration is guided by signaling through two receptor tyrosine kinases, PVR and EGFR, and blocking both receptors severely delays delamination (Duchek and Rørth, 2001; Duchek et al., 2001). To produce a guidance-deficient condition, we expressed DN forms of PVR and EGFR specifically in border cells. These cells’ front extensions during the wild-type delamination period were shorter and less persistent than those of wild-type cells (Fig. 4A,D,E, Movie 7). The guidance-deficient cells had normal locomotive behavior (Fig. 4C). However, although the guidance-deficient cells eventually moved sideways and delaminated, in most cases their route was less direct than that of wild-type cells (Fig. 4B, Movie 8). As the guidance-deficient cells approached the delamination point, their front extensions and movements were much like those observed in earlier stages of the period of analysis (Fig. 4C-E). We further found that the distance from the anterior tip of the egg chamber to the tip of the border-cell extension (defined as reach) was similar in wild-type and guidance-deficient mutants at the point of delamination (Fig. 4F). Wild-type and guidance-deficient border-cell clusters reached to approximately the same point before detaching completely, which suggests that the extension tip reached substrates it could grab hold of to aid in detaching the cluster, and that it is the cluster’s ability to reach such a substrate, rather than the actual length of the front extension, that is important for delamination. Consistent with this idea, the maximum length of the front extension did not correlate with the point of delamination, and the length of the front extension at the time of delamination varied more than the distance from the tip of the extension to the anterior tip of the egg chamber (Fig. S3). Locomotion and front extensions were similar in *shg*-mutant and guidance-deficient border cells (Fig. 3C,G,H, Fig. S2). However, *shg*-mutant border cells could not delaminate in the same time frame as guidance-deficient mutants, which might indicate that *shg*-mutant extensions were less able to grab onto of nurse cell substrates. Dai et al. recently revealed that a border cell’s preferential path is a juncture with multiple nurse cells (Dai et al., 2020). We found a hot spot close to the first multi-cell juncture with more than 3 nurse cells, suggesting that the preference for multi-cell junctures comes from not just more room, but also more membrane to grab hold of.

**Fig. 4.**
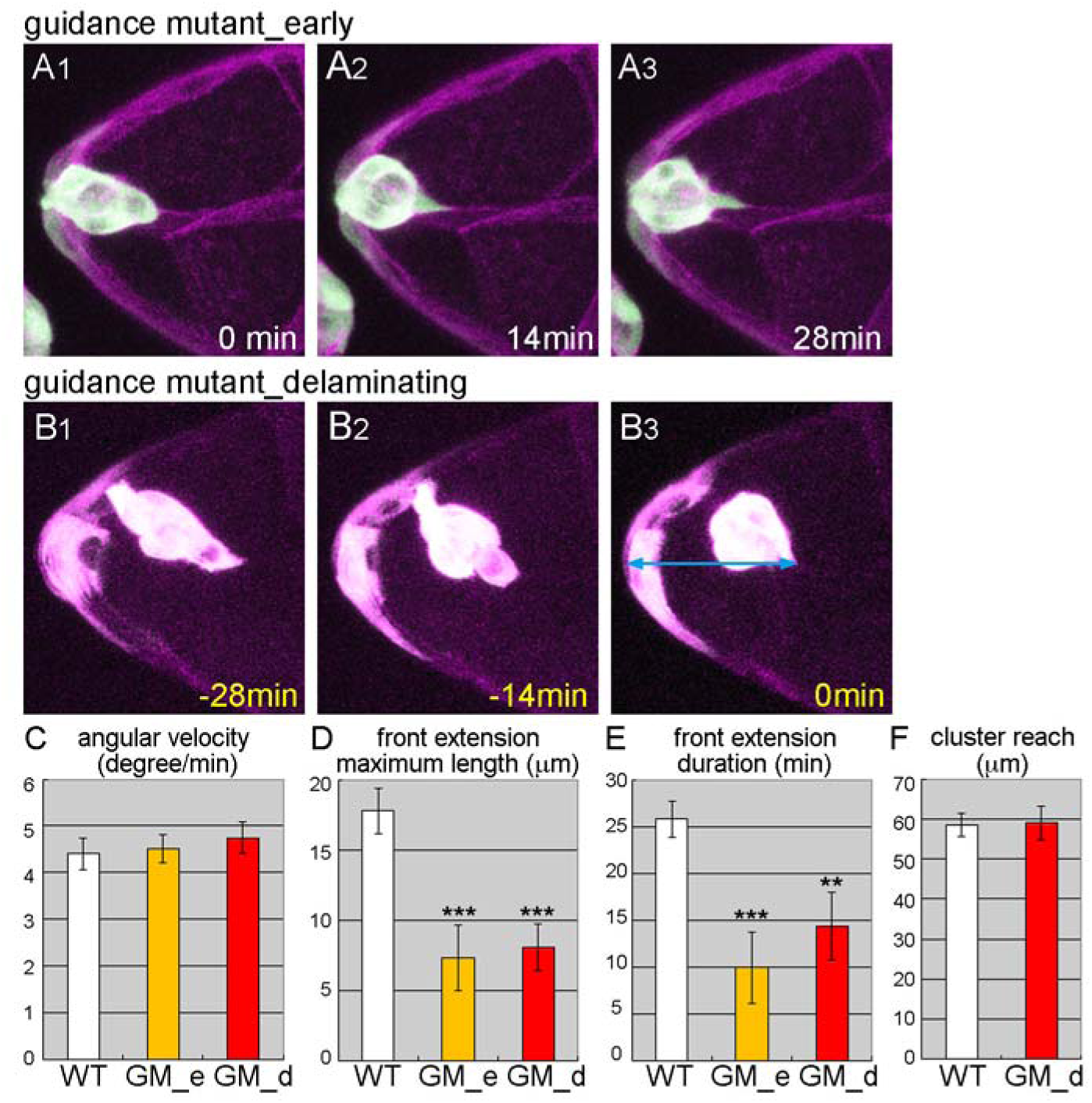
Invasive protrusion is required to reach a distance. (A,B) Z stacks from movies of guidance-deficient mutants in the wild-type (A), and the actual (B) delamination period. Light blue arrow indicates from the anterior end of the egg chamber to the reach of the border-cell cluster at delamination (B3). (C-E) The angular velocity (C), maximum length (D), and persistence (E) of front extensions of border-cell clusters in wild-type (WT), and guidance-deficient mutants in the wild-type (GM_e) and actual (GM_d) delamination periods. (C) 16≦n≦24, 8≦N≦12. (D,E) 8≦N≦14 (F) The distance from the anterior end of the egg chamber to the tip of the border-cell extension at the delamination point for wild-type (WT) and the guidance-deficient mutant (GM_d). 8≦N≦13. Error bars indicate SEM. ***, p<0.001, **, p<0.01. Time shown at the lower left indicates time elapsed from the start of the movie (white) and time prior to delamination (yellow). For all images, anterior is left.

### Motility is not a prerequisite for invasiveness

E-cad induces invasive extensions in *slbo*-mutant border cells, which normally have no detectable extensions (Fig. 3B,G,H). We examined whether E-cad could induce invasive extensions in JAK/STAT-inhibited border cells, which lack locomotion. Under normal conditions, border cells appear to send out extensions after acquiring motility (Fig. 1A). To our surprise, forced E-cad expression induced JAK/STAT-inhibited follicle-like cells to send out front extensions, without changing the morphology of other follicle cells in most cases (Fig. 5A, Movie 9), and the length and duration of these front extensions were very like wild-type extensions (Fig. 5D,E). Thus, non-motile cells were able to form extensions, confirming that motility and invasiveness are regulated independently. We investigated this possibility further by overexpressing Slbo in JAK/STAT-inhibited border cells. JAK/STAT-inhibited follicle-like cells with forced Slbo expression produced front extensions, as did those with E-cad (Fig. 5B, Movie 10), but other border cells were unaffected in shape and were less motile than wild-type (Fig. 5C), supporting the idea that invasive extension does not require cell motility. These results also ruled out the possibility that residual Slbo activity conferred motility to the *slbo^1310^* mutant. Both E-cad and Slbo overexpression induced extensions that were not sufficiently invasive for successful delamination.. Extensions induced by E-cad were very thin and grew without ever retracting, suggesting that they lacked machineries to produce traction forces (Fig. 5A4). In contrast, extensions induced by Slbo were thicker and retractable, suggesting that other downstream components of Slbo serve to make the extensions functional. This conclusion is consistent with the results of rescue experiments in *slbo* mutants (Fig. 5B4,3D).

**Fig. 5.**
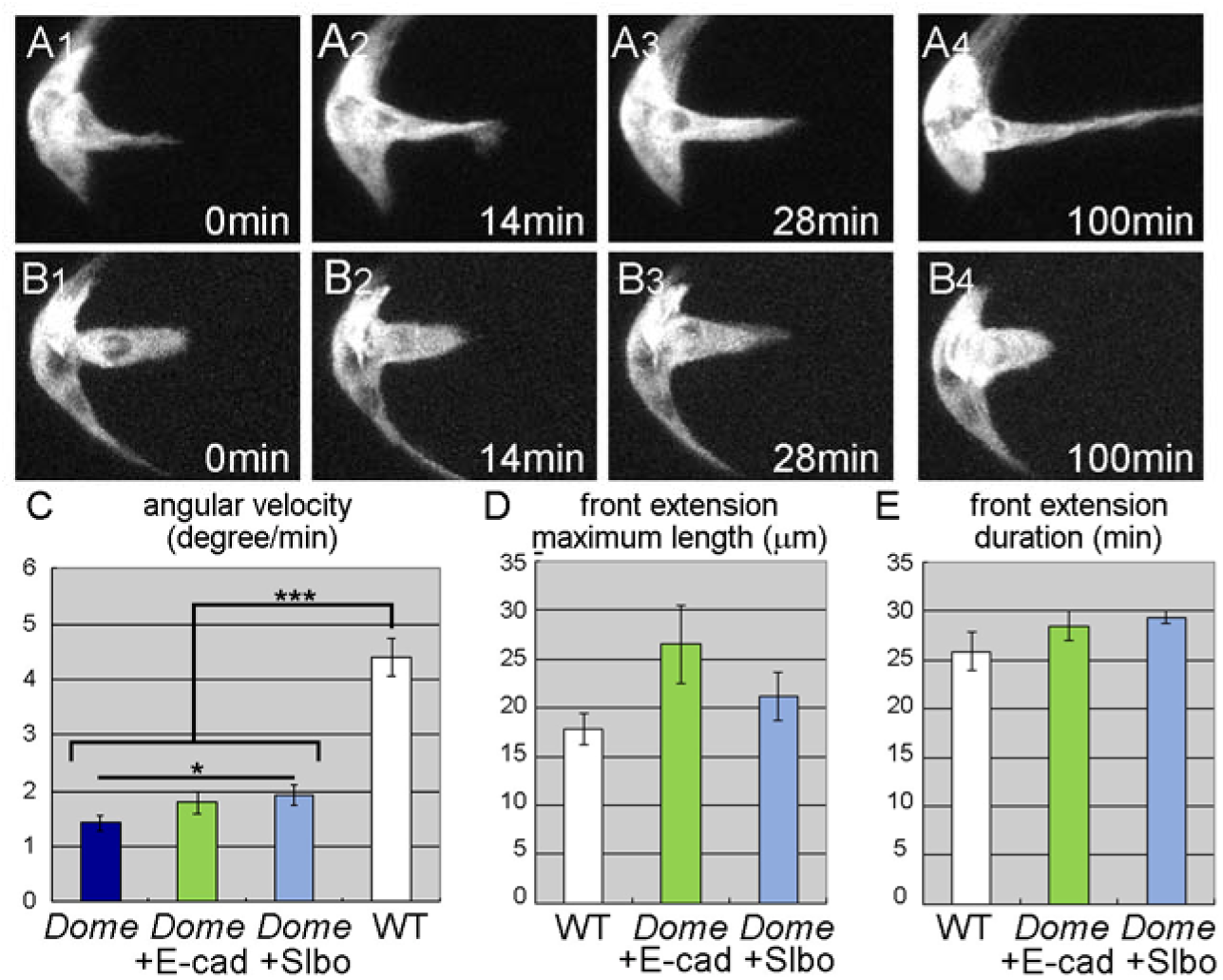
Motility is not a prerequisite for invasiveness. (A,B) Z stacks from movies of border cells expressing DN-Dome with E-cad (A) and Slbo (B). E-cad and Slbo induced front extensions in JAK/STAT-inhibited border cells. (C-E) Angular velocity and (C) the maximum length (D) and duration of front extensions in wild-type (WT) or DN-Dome expressing (*Dome*) border-cell clusters with either E-cad (*Dome*+E-cad) or Slbo (*Dome*+Slbo) expression. (C) 16≦n≦24, 8≦ N≦12. (D,E) 6≦N≦14. Error bars indicate SEM. ***, p<0.001, **, p<0.01. Time elapsed from the start of the movie is indicated at the bottom right. For all images, anterior is left.

## Discussion

Epithelial cells undergo a series of cellular events as they approach delamination. Cells de-adhere from neighboring cells, begin moving actively as individual cells, send protrusions toward the tissue they will move into, and then detach from the original cell layer. We showed that motility and invasiveness are acquired independently in delaminating *Drosophila* border cells. Border cells are differentiated by JAK/STAT signaling. Downstream of JAK/STAT signaling, the transcription factor Slbo regulates border cell invasiveness, while motility is regulated by other downstream factors. In models of cancer metastasis, cell proliferation and migratory behavior are thought to be genetically distinct. However, cell motility and invasiveness are not often considered separately (Cagan et al., 2019; Stuelten et al., 2018). Here, we revealed that motility and invasiveness are regulated independently in delamination. In wild-type border cells, individual cells begin to jiggle slightly, develop small spikey extensions, and eventually form a round cluster that sends out a single large extension (Fig. 1A). Thus, it might seem that cells first acquire motility and then invade the path of migration. However, our results showed that cell motility and invasiveness are acquired independently; thus, cell motility is not a prerequisite for invasiveness. This was confirmed when forced Slbo expression in JAK/STAT-inhibited border cells rescued invasive extensions without affecting cell motility (Fig. 5B-E). Although guidance-defective border cell clusters produced extensions that were smaller and less persistent than wild-type, these clusters eventually delaminated (Fig. 4D,E) by sideways movements (Fig. 4B). Thus, the locomotive and invasive behaviors of border cells cooperatively contribute to delamination.

Acquiring motility involves actin cytoskeleton regulators, since inhibiting actin polymerization diminished border cell motility (Fig. 2E,G). Border cells retain apicobasal polarity during delamination (Pinheiro and Montell, 2004; Wang et al., 2018). Thus, border cells may become motile and form clusters without any major changes in the distribution of proteins involved in apicobasal polarity. Changes in adhesion between border cells and neighboring follicle cells may affect the shape and formation of border-cell clusters. Some of these properties might be regulated downstream of JAK/STAT signaling by transcription factors other than Slbo. The few downstream targets of JAK/STAT signaling identified in border cells include Apontic and Socs36, both of which modulate JAK/STAT signaling itself (Monahan and Starz-Gaiano, 2013; Starz-Gaiano et al., 2008).

The acquisition of invasiveness involves E-cad downstream of Slbo. E-cad is required in both border cells and nurse cells for border cell migration (Mathieu et al., 2007). E-cad is required for border cell protrusions, which are formed by homophilic interaction with E-cad on the nurse cells. Although it might appear that E-cad upregulation in border cells is involved in cluster integrity, E-cad upregulation is transcriptionally regulated by Slbo (Mathieu et al., 2007), and a lack of Slbo only affects the invasiveness of the border-cell cluster (Fig. 2C). Furthermore, forced expression of E-cad in the JAK/STAT-inhibited border-cell region did not restore border-cell clusters to a round shape (Fig. 5A). Thus, E-cad upregulation in border cells contributes solely to invasiveness. Cluster extensions induced by forced Slbo expression were more functional than those induced by E-cad (Fig. 5B), suggesting that other factors are involved in the formation, maintenance, and retraction of invasive extensions. These factors likely include actin cytoskeleton regulators. Extensive searches for Slbo targets (Borghese et al., 2006; Wang et al., 2006) have identified some actin-associated proteins, including Myosin VI, encoded by *jargur*, and Fascin, encoded by *singed* (Borghese et al., 2006; Geisbrecht and Montell, 2002). Myosin VI stabilizes E-cad in border cells, and Fascin regulates stiffness in nearby nurse cells (Geisbrecht and Montell, 2002; Lamb et al., 2020; Lamb et al., 2021). To generate the necessary traction forces for delamination, it is likely that more actin-associated proteins are involved in border-cell extension, or that proteins that have already been identified play additional roles. To clarify their contributions, reconstruction experiments should be conducted using multiple downstream genes.

## Materials and methods

### Fly strains

All flies were raised at 25 °C with standard media. We used the following fly strains: *slbo*-gal4 (Rørth et al., 1998), *slbo^1310^* (Montell et al., 1992), *slbo^e7b^* (null, Rørth, 1994), *shg^PB4354^* (Mathieu et al., 2007), UAS-*Dome*^Δ^*^CYT^* (Silver and Montell, 2001), UAS-*shg* (Mathieu et al., 2007), UAS-*slbo* (Rørth et al., 2000), UAS-DN-PVR, UAS-DN-EGFR (Duchek and Rørth, 2001), and UAS-10×GFP (Poukkula et al., 2011). We used the following genotypes for reconstruction experiments: *slbo^1310^*/ *slbo^1310^*; UAS-*shg* / *slbo*-gal4, UAS-10×GFP, UAS-*shg*, UAS-*Dome*^Δ^*^CYT^* / *slbo*-gal4, UAS-10×GFP, and UAS-*slbo*, UAS-*Dome*^Δ^*^CYT^* / *slbo*–gal4, UAS-10×GFP.

### Live imaging and drug treatment

Live imaging was as previously described (Bianco et al., 2007). Newly emerged females of the appropriate genotypes were collected and maintained one day on dry yeast followed by one day on fresh yeast. Ovaries were taken in Schneider’s medium (Gibco, 21720) containing 5 μg/ml of Insulin (Sigma-Aldrich, 19278). Egg chambers were dissected out in the media and moved into cover-glass chambers (Lab-Tek,155411) coated with 0.1 mg/ml poly-L-Lysine (Sigma-Aldrich, P7405) using truncated pipette tips. We added 250μm of imaging media, consisting of Schneider’s medium containing 2.5% Fetal Bovine Serum (heat-inactivated, Biowest), 5 μg/ml Insulin, 2 mg/ml Trehalose (Sigma-Aldrich, 90208), 5 μM Methoprene (Sigma-Aldrich, 33375), 1 μg/ml 20-hydroxyecdysone (Sigma-Aldrich, H5142), and 9 μM FM4-64 (Invitrogen, T13320). Images were acquired by confocal microscopy (SP5, Leica) with a 63× objective; sections were 2.5 μm apart and covered the cluster every 2 min for 2 h. For quality control of the movies, we used transmission images to see whether the growth and cellular integrity of the egg chamber were maintained for 2 h. To inhibit actin polymerization, CytochalasinD (Sigma, C8273) was added to the imaging media at a final concentration of 25 μM.

### Image analysis and quantification

For egg chamber staging, we selected transmission images of single Z slices showing the maximum egg chamber area, and measured the width and height of the egg chamber and the oocyte. We selected oocyte length as a marker for the wild-type delamination period, corresponding to an oocyte length of 50–70 microns. We analyzed movies of wild-type *Drosophila* for 30 min prior to the delamination point; for mutants, we analyzed the 30 min period starting with the first frame in the wild-type delamination period (assessed according to oocyte width). For guidance-receptor mutants, we analyzed both time periods. To analyze single-cell movement, we tracked individual cells using nuclear position, marked by absence of GFP expression. We tracked the center of the anterior pole cells as the cluster center. We used an Image J macro to measure changes in the angle of border cells relative to the cluster center, and calculated the angular velocity. We analyzed front extensions using custom macros slightly modified from Poukkula et al. (2011). After making binarized images, we manually deleted stretched cells that linked border-cell clusters, then analyzed extensions as described previously (Poukkula et al., 2011). We could not use the same method for rescue experiments for JAK/STAT inhibition because the border cells did not form a round cluster, so we separated extensions by drawing a line where the extension base touched other border cells and measured the length and duration of the extensions. Statistical analyses were performed using Welch’s or Student’s t tests, depending on the results of verification of standard deviation by F test. We calculated two-sided p values.

## Acknowledgements

We would like to thank Pernille Rørth and the Institute of Molecular and Cell Biology in Singapore for their support. This work was supported by Sumitomo, Futaba, and Takeda Science Foundations.

## Movies

Movie 1

Delaminating wild-type *Drosophila* border cells form a cluster of jiggling cells. The cluster extends a protrusion in the direction of future migration, and the tip finds an attachment point. The cell cluster detaches from the epithelial layer and follows the protrusion as it contracts. Elapsed time (minutes) shows at lower right.

Movie 2

JAK/STAT-inhibited follicle cells failed to differentiate into border cells, move into clusters, or extend protrusions in the period corresponding to wild-type delamination. JAK/STAT signaling was inhibited with DOME lacking a cytoplasmic domain. Elapsed time shows at the lower right.

Movie 3

*slbo*-mutant border cells formed clusters and moved within the clusters, but did not form protrusions or delaminate during the period when wild-type cells would delaminate. Elapsed time shows at the lower right.

Movie 4

*slbo-*mutant border cells formed clusters but lost motility when treated with CytochalasinD, which inhibits actin polymerization, and failed to delaminate during the period of wild-type delamination period. Elapsed time shows at the lower right.

Movie 5

*shg*-mutant border cells formed clusters of motile cells and sent out short extensions, but did not delaminate during the wild-type delamination period. Elapsed time shows at the lower right.

Movie 6

*slbo*-mutant border cells with E-cad expression formed front extensions but did not delaminate during the wild-type delamination period. Elapsed time shows at the lower right.

Movie 7

Guidance-deficient border cells had normal motion but formed front extensions that were shorter and less persistent than wild-type extensions.

Movie 8

Guidance-deficient border cells delaminated successfully. Elapsed time shows at the lower right.

Movie 9

E-cad expression did rescued invasive extensions in JAK/STAT-inhibited border cells did not rescue cell movement. Elapsed time shows at the lower right.

Movie 10

Slbo overexpression induced front extensions in JAK/STAT-inhibited follicle-like cells, but the cells failed to delaminate. Elapsed time shows at the lower right.

---

**Fig. S1.**
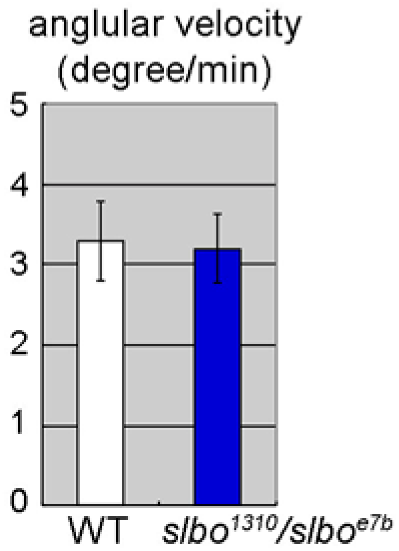
The angular velocity of wild-type (WT) and *slbo^1310^/slbo^e7b^* border cells. Angular velocities were comparable in these conditions. n=10, N=5. Error bars indicate SEM.

**Fig. S2.**
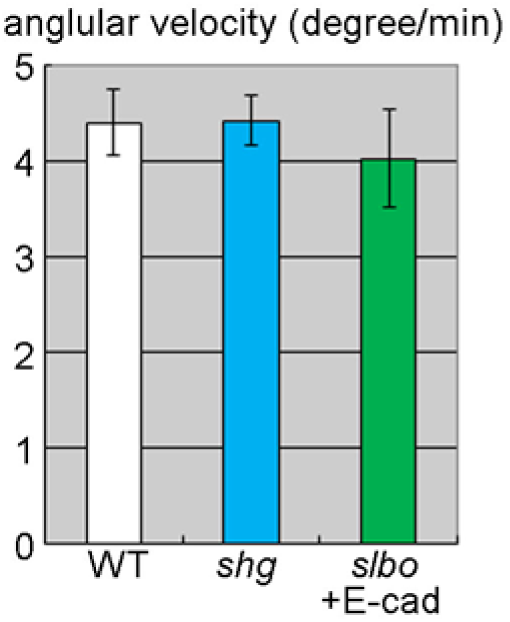
Angular velocity for the following border cells: wild-type (WT), *shg*-mutant (*shg*), and *slbo*-mutant with E-cad expression (*slbo*+E-cad). Angular velocities were comparable in these conditions. 10≦n≦26, 5≦N≦13. Error bars indicate SEM.

**Fig. S3.**
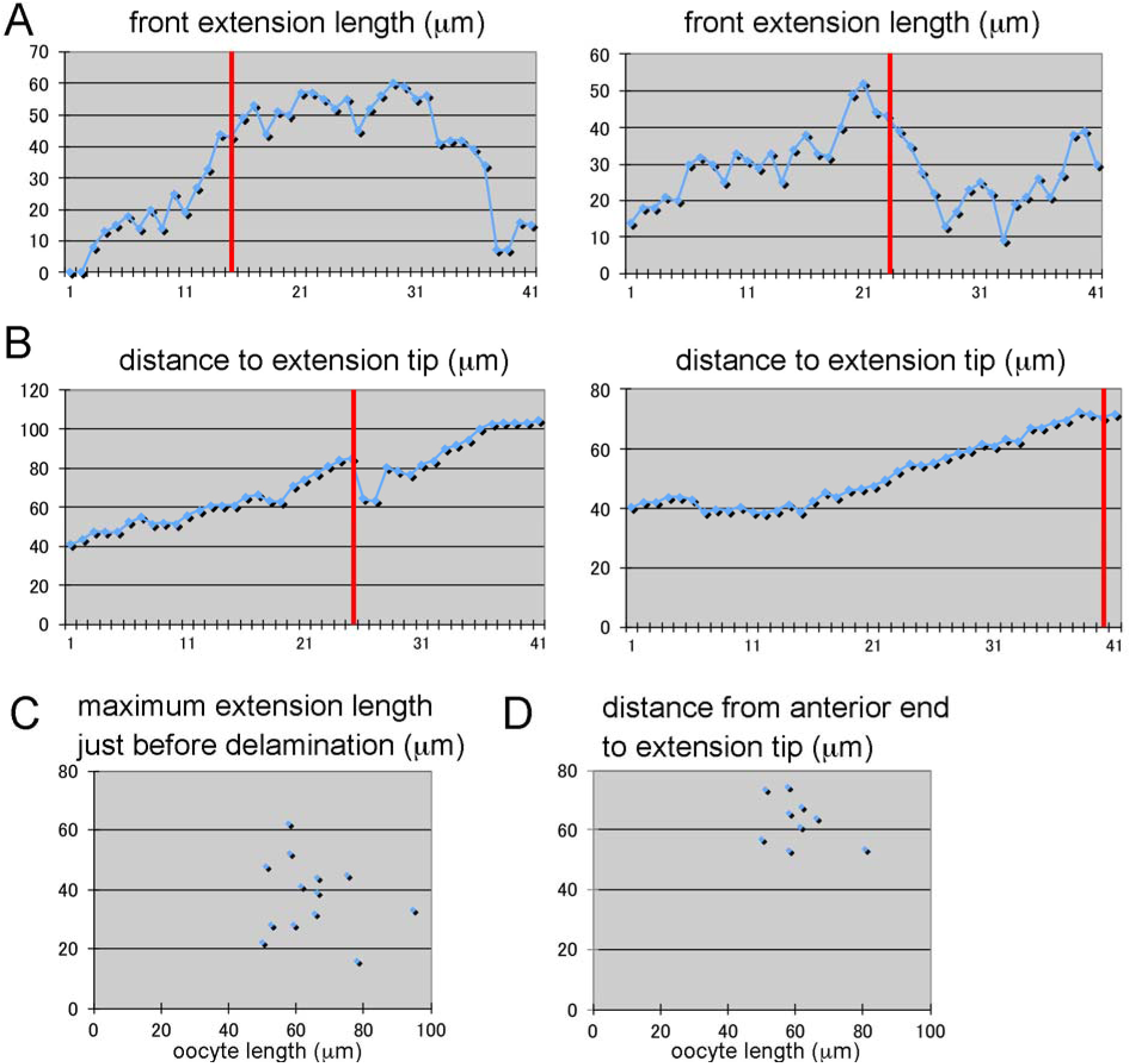
(A,B) Examples of changes in front extension length (A) and in cluster reach (the distance between the anterior end of the egg-chamber and the extension tip) (B) during delamination. X axis: time frame. Red lines: delamination point. Front extension length varied quite a bit, while cluster reach tended to increase over time. The maximum extension length did not correlate with the delamination point. (C,D) The maximum extension length (C) and the cluster reach (D) just before delamination. Length varied more than reach.

